# Distinct roles of spinal commissural interneurons in transmission of contralateral sensory information

**DOI:** 10.1101/2023.02.16.528842

**Authors:** Olivier D. Laflamme, Sergey N. Markin, Rachel Banks, Ying Zhang, Simon M. Danner, Turgay Akay

## Abstract

Crossed reflexes (CR) are mediated by commissural pathways transmitting sensory information to the contralateral side of the body, but the underlying network is not fully understood. Commissural pathways coordinating the activities of spinal locomotor circuits during locomotion have been characterized in mice, but their relationship to CR is unknown. We show the involvement of two genetically distinct groups of commissural interneurons (CINs) described in mice, V0 and V3 CINs, in the CR pathways. Our data suggest that the exclusively excitatory V3 CINs are directly involved in the excitatory CR, and show that they are essential for the inhibitory CR. In contrast, the V0 CINs, a population that includes excitatory and inhibitory CINs, are not directly involved in excitatory or inhibitory CRs but down-regulate the inhibitory CR. Our data provide insights into the spinal circuitry underlying CR in mice, describing the roles of V0 and V3 CINs in CR.

## MAIN

Crossed reflexes are motor responses of one limb to somatosensory stimuli applied to the contralateral limb. These crossed reflex pathways has been shown to be excitatory, initiating motor response on the contralateral leg, as well as inhibitory, suppressing motor activation on the contralateral leg ^1–7^. The spinal circuitry that mediates crossed reflexes has been investigated in detail leading to the identification of multiple commissural interneurons mediating sensory information to the contralateral side of the spinal cord ^1–7^. Moreover, it has been shown that crossed reflexes can be initiated by simulating contralateral cutaneous ^7,8^ as well as proprioceptive ^1–7^ afferent fibers. Although, many details of these pathways have been described, the network that controls these crossed reflexes is obscure.

It has been argued that the significance of crossed reflex pathways lies in their importance for interlimb coordination, and to maintain stability while standing and during locomotion, for example during stumbling correction ^9–11^. These abilities are compromised following various motor disorders ^12–15^ as well as in the elderly ^16^. Indeed, impairment of crossed reflexes has been shown in individuals with stroke ^17,18^. Therefore, understanding the spinal circuitry that underlies crossed reflexes are important not only for basic science research to understand function of the central nervous system, but also for medical research.

Advances in mouse developmental genetics and genetic engineering led to the description of spinal neuronal circuitry that coordinates the movement of the left and right hind limbs during locomotion at different speeds ^9,19^. Two main cardinal groups of commissural interneurons, the V0 and the V3, have been shown to emerge from two ventrally located progenitor domains, the p0 and the p3, of the embryonic spinal cord ^20–22^. While the V0 further diverges into dorsal inhibitory CINs (V0_d_) and ventral excitatory CINs (V0_v_) ^20^, the V3 CINs, also diverging into three subgroups that are exclusively excitatory^23,24^. Previous research has shown that the removal of the V0 CINs from the spinal network using mouse genetics severely disrupts left–right alternation during locomotion in a speed-dependent manner ^25,26^. Furthermore, the subgroups of the V3 and V0 CINs has been shown to play different roles in control of distinct behaviors ^23,27^. In contrast, genetically silencing V3 CIN output does not eliminate left–right alternation during locomotion, but it compromises the stability of interlimb coordination ^21,28^. It was suggested that the V3 CINs might have a role in synchronization of motor activity in the left and right limbs seen in galloping or bounding ^26,28–30^. Given the importance of V0 and V3 CINs for left–right coordination during locomotion, we tested the hypothesis that the V0 and V3 CINs mediate crossed reflex pathways.

To test our hypothesis, we investigated the involvement of V0 and V3 commissural interneurons in the crossed reflexes by combining mouse genetics with the in vivo methodology we recently developed^31^. Our data provide evidence that: 1) The V3 CINs, although not necessary for the alternating movement of the left and right legs of the same segment during locomotion are involved in excitatory crossed reflexes and are necessary for the inhibitory crossed reflexes. 2) Although necessary for left–right coordination during locomotion, the V0 CINs are not necessary for crossed reflexes, but they have a modulatory influence on the crossed reflex actions. Taken together, the findings demonstrate the basic structure of spinal commissural circuits underlying crossed reflexes in mice. Furthermore, our data suggest that the genetically defined commissural interneurons, V0 and the V3 CINs, have distinct function in crossed reflexes. While V3s are directly involved mediating crossed reflex responses, the V0 CINs exerts rather a modulatory function.

## RESULTS

### Involvement of V0 and V3 CINs in the excitatory crossed reflex pathways

To measure the crossed reflex responses of different leg muscles to afferent activation on the contralateral leg, we recorded electromyogram (EMG) activities from multiple muscles of the right hind limb while we electrically stimulated either the tibial nerve or the sural nerve of the left hind limb (**Fig.1a**). Stimulation of the nerves was carried out either at low (1.2xT) or high (5xT) threshold intensity necessary to elicit local response in the left gastrocnemius muscle (Gs). Based on previous research ^32,33^, 1.2xT stimulation of the tibial nerve will activate mostly proprioceptive afferent fiber of groups Ia and Ib that innervate the triceps surae muscles and the 5xT stimulation will additionally activate the group II afferents that include proprioceptive group II afferents from muscles and cutaneous low threshold mechanoreceptors. Stimulation of the sural nerves is likely to activate cutaneous afferents of large caliber at 1.2xT and of intermediate to large caliber at 5xT. However, it has to be considered that in rodents the sural nerve also harbors minor motor fibers that innervate the flexor digiti minimi muscle ^33,34^. Therefore, minimal efferent or proprioceptive contribution during sural nerve stimulation cannot be excluded. These experiments were carried out on wild type mice and in mutant mice where either V0 CINs were killed (V0^kill^) ^25^, or the synaptic output of V3 CINs was silenced (V3^off^) ^24^.

**Figure 1:**
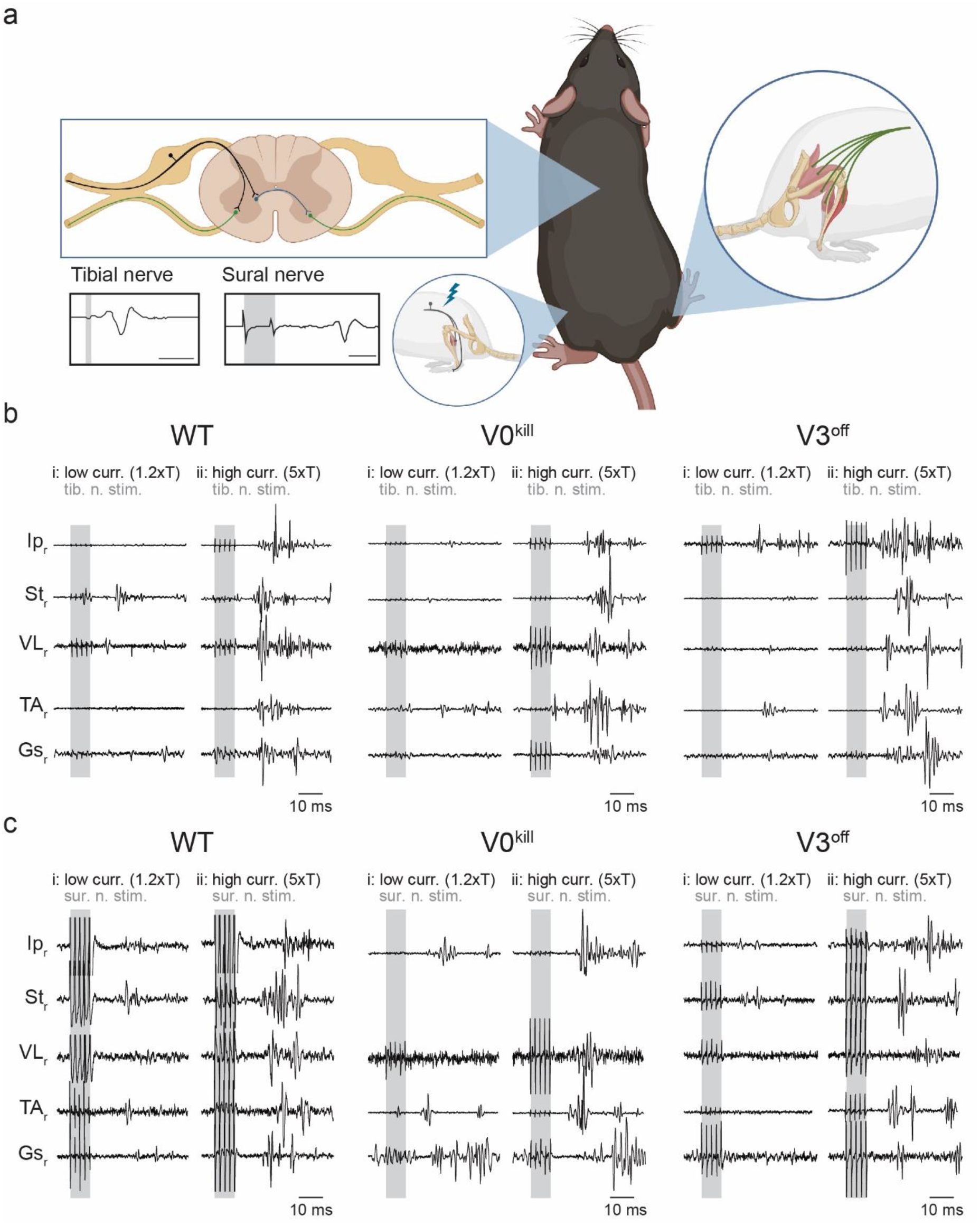
Crossed reflex responses in wild type, V0^kill^ and V3^off^ mice. **a,** Schematic of the experimental setup. Nerve cuff electrodes to the left tibial or sural nerve along with one EMG recording electrode into the left Gs were implanted to determine the threshold current to activate local reflex. In addition, EMG recording electrodes to five muscles of the left leg were implanted to record crossed reflex responses. Schematic was created with BioRender.com (agreement number: QQ250Q4M08). **b,** EMG responses of all recorded muscles of the right leg to left tibial nerve stimulation at 1.2xT (i) and 5xT (ii) in wild type (WT), V0^kill^ and V3^off^ mice. Shaded areas indicate nerve stimulation. **c,** Same as in **b**, but for responses to left sural nerve stimulation.

In wild type mice, stimulation of the left tibial nerve or sural nerve at 1.2xT or 5xT activated all recorded muscles (**Fig. 1b and 1c**). Visual comparison of average rectified EMG activities from all muscles in V0^kill^ and V3^off^ mice indicated that crossed reflex responses could be elicited in the absence of V0 and V3 CINs. Furthermore, the overall activity pattern qualitatively resembled the responses recorded in wild type mice when the left tibial (**Fig. 2a**) or sural (**Fig. 2b**) nerve was stimulated. Next, we sought to quantify the muscle responses in all mice, all muscles, and after both nerve stimulations. To do this, we measured the area underneath the average rectified EMG traces within a 12–50 msec time window after the stimulation onset (**Fig. 2c and 2d**). With increasing intensity from 1.2xT to 5xT responses increased (ratio=1.84±0.96, p<0.0001); the extent of the increase did not differ between WT, V0^kill^ and V3^off^ mice (interaction effect: F_2,283_=2.286, p=0.103). There was significant effect of mouse line (F_2,283_=7.750, p=0.0005): across muscles, nerves, and stimulation intensities, V3^off^ mice exhibited significantly smaller muscle activities than V0^kill^ mice (ratio=0.687±0.078, p=0.0030) and their wild-type counterparts (ratio=0.663±0.077, p=0.0013) (**Fig. 2e**). All second and third order interaction effects involving mouse type and nerve and/or stimulation intensity were not significant (all p>0.1), indicating that the reduction of amplitudes in V3^off^ mice compared to WT and V0^kill^ mice was present independently of stimulated nerve and stimulation intensity. Yet, the interaction between mouse type (WT, V0^kill^, and V3^off^) and muscle was significant (F_4,283_=8.370, p<0.0001) indicating that the effect of mouse type was different between muscle groups. Indeed, responses in V3^off^ mice were significantly smaller than in WT mice only in Ip (ratio=0.574±0.087, p=0.0009), VL (ratio=0.588±0.091, p=0.0019), and St (ratio=0.399±0.084, p=0.0001); and smaller than in V0^kill^ mice in St (ratio=0.457±0.09, p=0.0003) and TA (ratio=0.625±0.12, p=0.0380). These data suggested that V3 CINs, but not the V0 CINs, might have a role, albeit minor, in mediating excitatory crossed reflex responses.

**Figure 2:**
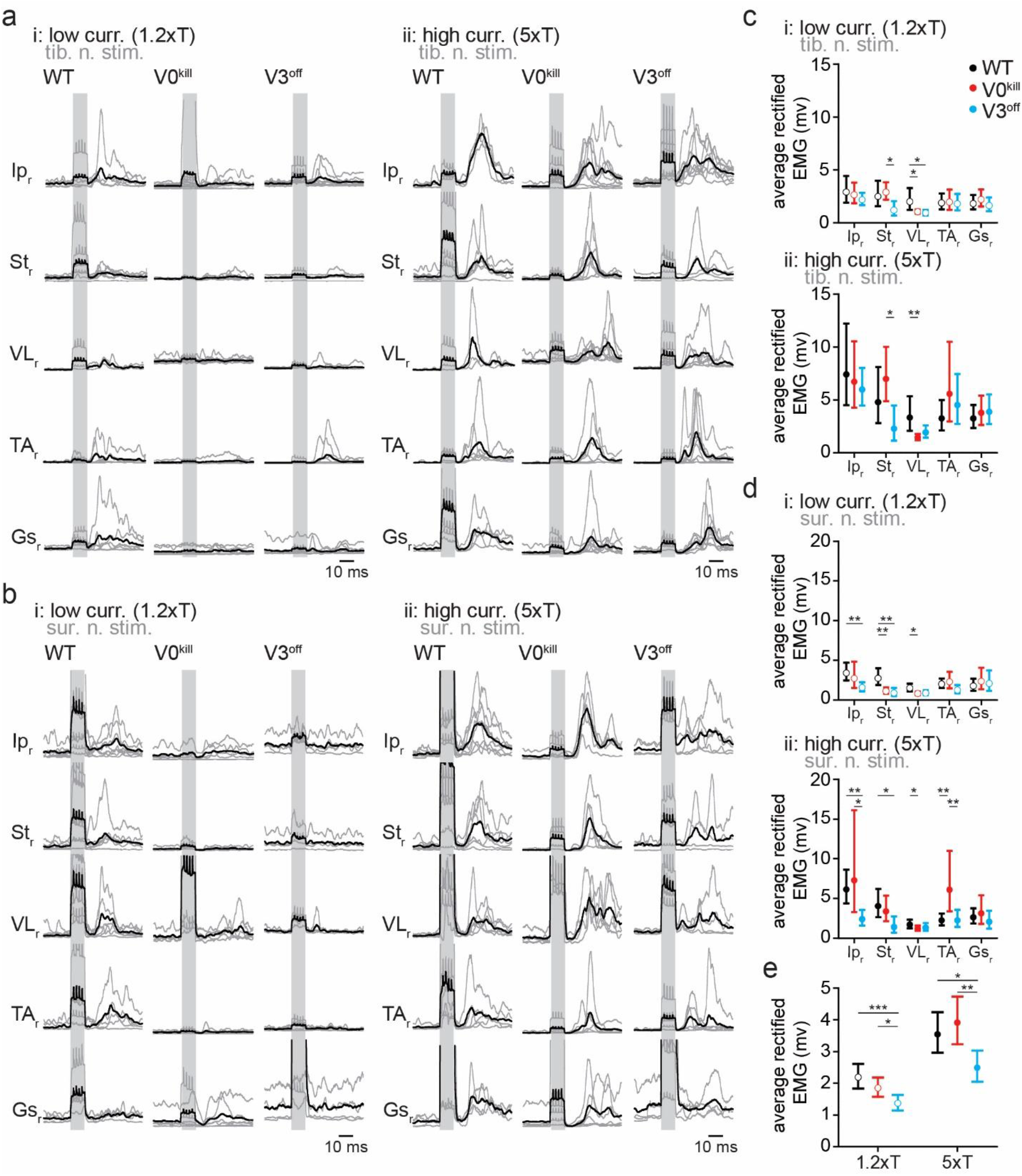
Average EMG responses of muscles to contralateral nerve stimulation. **a,** Average traces of rectified and filtered EMG activities from right leg muscles as a response to left tibial nerve stimulation at 1.2xT (i) and 5xT (ii) in wild type, V0^kill^ and V3^off^ mice. The gray lines are averages from individual animals, whereas the black lines are averages across all animals. The shaded backgrounds indicate the time when the left nerves were stimulated. **b,** Same as in a but for sural nerve stimulation. **c,** Graphs showing the average area underneath the average rectified EMG traces from 12–50 msec delays (the delay was chosen to exclude influence from stimulation artifacts) from the stimulation onset for 1.2xT (top) and 5xT (bottom) stimulation of the tibial nerve. The error bars indicate the 95% confidence intervals. **d,** Same as in c but for sural nerve stimulation. **e,** Graphs illustrating the combined averages from all EMG traces for 1.2xT and 5xT stimulation of the tibial and the sural nerves combined. *: p<0.05, **: p<0.01, ***: p<0.001.

To further elaborate on this, we next investigated the likelihood of eliciting a muscle response with stimulation of the left tibial nerve and sural nerve (**Fig. 3**). This was done by visually inspecting EMG traces immediately after nerve stimulation. The relative frequency of occurrence of muscle activity after nerve stimulation was calculated as the number of nerve stimuli that were followed by a muscle response over the total number of stimulations. Across muscles, stimulated nerves, and mouse types, the frequency of muscle activity in response to stimulation was significantly higher at 5xT than at 1.2xT [F_1,258_=396.093, odds ratio (OR)=6.94±0.676, p<0.0001, **Fig. 3a**]. While there wasn’t a significant difference of the frequency of responses between WT, V0^kill^, and V3^off^ mice (F_2,258_=0.364, p=0.3586), the interaction effect with stimulation intensity was significant (F_2,258_=21.745, p<0.0001), indicating that the influence of stimulation intensity differed between the mouse types (**Fig. 3b**). Indeed, at 1.2xT significantly fewer responses were recorded in V3^off^ mice than in WT (OR=0.458±0.130, p=0.0171) but not V0^kill^ mice (OR=0.680±0.191, p=0.3577). Furthermore, at 5xT significantly more responses were recorded in V0^kill^ mice than WT mice (OR=2.449±0.721, p=0.0073). The lack of a significant three-way interaction effect with nerve type (F_8,258_=0.778, p=0.355) indicates that this effect is not influenced by whether tibial (**Fig. 3c**) or sural nerve (**Fig. 3d**) was stimulated. In summary, both, the EMG activities in response to the left nerve stimulation and the reliability of these responses at low stimulation intensities, were lower in V3^off^ mice than in their wild-type counterparts. Together, these data suggest that V3 CINs but not V0 CINs have a minor contribution to the excitatory components of the crossed reflexes.

**Figure 3:**
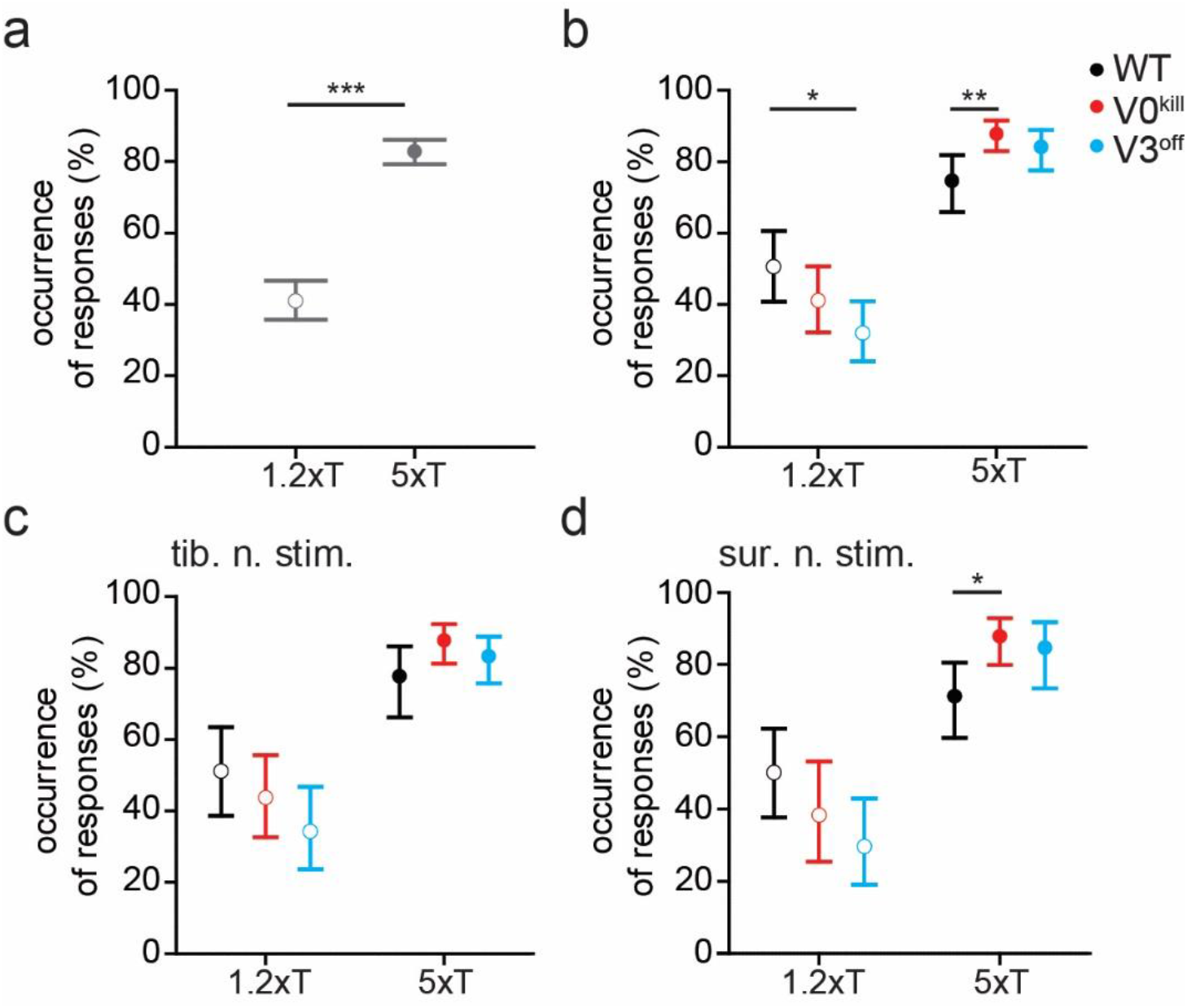
The probability of muscle activation does not change in the presence or absence of V0 or the V3 CINs. **a,** Occurrence of observed responses across all recorded muscle in all three groups of mice when either nerve tibial or sural nerve was stimulated either at 1.2xT or 5xT, indicating higher rate of muscle activity was observed with higher stimulation strength. **b,** Increasing the stimulation strength from 1.2xT to 5xT increases the occurrence of observation in muscles similarly in all three mouse groups. Data from both nerve stimulations are combined in this graph. **c** and **d,** The effect of nerve stimulation at 1.2xT or 5xT is regardless of whether the tibial nerve (**c**) or the sural nerve (**d**) is stimulated. *: p<0.05, **: p<0.01, ***: p<0.001.

Next, we asked whether there are temporal differences in the muscle activity patterns during crossed reflex responses between the three groups of mice. To do this, we detected the delays from the stimulation onset to the on and offsets of muscle activity responses to the tibial or sural nerve stimulations at 1.2xT and 5xT (**Fig. 4**). The fixed effect of mouse type was significant (F_2,225_=4.921, p=0.0081), indicating that across stimulated nerves, stimulation intensities, and muscles, onset latencies differed between WT, V3^off^, and V0^kill^ mice (**Fig. 4a**); V0^kill^ mice exhibited significantly longer latencies than WT mice (ratio=1.12±0.0405, p=0.0063). Longer latencies could indicate a stronger inhibitory crossed reflex component in V0^kill^ mice compared to WT mice. Across mice, stimulated nerves, and muscles, onset latencies were significantly higher at 5xT stimulation than at 1.2xT stimulation (F_2,225_=10.765, p=0.0012; **Fig. 4b**). This latency increase with stimulation intensity is in accordance with our previous data ^31^ and suggests the presence of inhibition preceding the crossed-reflex response. Furthermore, there was a significant interaction effect between mouse type, nerve, and stimulation intensity (F_2,225_=8.508, p=0.0003), suggesting that the effect of stimulation intensity on the onset latencies differed between WT, V3^off^, and V0^kill^ mice and stimulated nerve (**Fig. 4c**). Indeed, this increase of latency with stimulation intensity was only significant for WT mice (ratio=1.239±0.041, p<0.0001) and V0^kill^ (ratio=1.102±0.046, p=0.0213) with tibial nerve stimulation; with sural nerve stimulation there was no significant change in latency between 1.2xT and 5xT in neither WT, V3^off^, or V0^kill^ mice (WT: ratio=0.991±0.0326, p=0.7922; V3^off^: ratio=1.09±0.0666, p=0.1618; V0^kill^: ratio=1.029±0.0537, p=0.5805); and even with tibial nerve stimulation V3^off^ mice did not show a change of onset latencies between stimulation intensities (ratio=0.948±0.0421, p=0.2318). This increase of onset latency was previously shown to be likely the result of an inhibitory crossed reflex pathway ^31^. In this case the data would suggest that the inhibitory component of the crossed reflex was compromised in V3^off^ mice.

**Figure 4:**
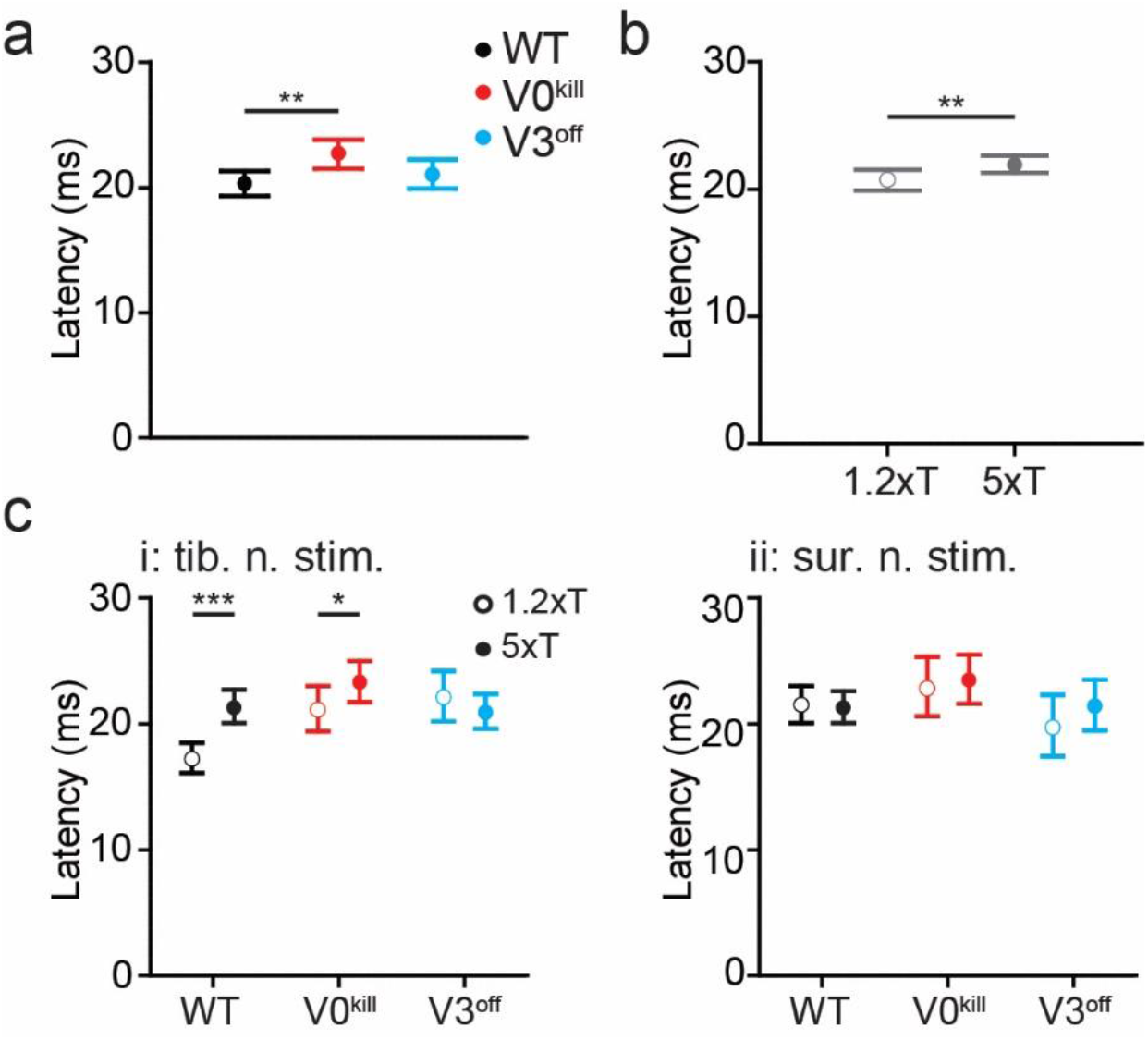
Temporal differences in muscle activation pattern in wild type, V0^kill^, and V3^off^ mice. **a,** In general, the latencies of muscle activation was larger in V0^kill^ mice than wild type and V3^off^ mice. **b,** Increasing the stimulation strength from 1.2xT to 5xT caused an increase in delay across all animal groups. Data from both nerve stimulations are combined in this graph. **c**, The stimulation strength dependent increase in latency shown in b was clearly present in wild type and V0^kill^ mice but not in V3^off^ mice, when tibial nerve was stimulated (i). The delays did not show any strength dependent change in any mouse groups when sural nerve was stimulated (ii). *: p<0.05, **: p<0.01, ***: p<0.001.

Our observations presented above suggests that V3 CINs contribute to the excitatory crossed reflexes. First, the overall EMG activity (**Fig. 2e**) as a response to the left nerve stimulation is lower in amplitude in V3^off^ mice compared to wild type and V0^kill^ mice. Second, the decreased occurrences of reflex responses at low current stimulation (**Fig. 3c** and **3d**) in V3^off^ mice following the left nerve stimulation is lower in V3^off^ mice than the overall activity in WT and V0^kill^ mice. No indication for V0 CIN contribution to the excitatory crossed reflex action could be detected. This suggests the V3 but not V0 CINs are likely to be part of the excitatory crossed reflex pathway.

### Involvement of V0 and V3 CINs in the inhibitory crossed reflex pathways

Next, we aimed to investigate the contribution of the V0 and the V3 CINs to the inhibitory crossed reflexes, using the double nerve stimulation technique we recently developed ^31^. To do this, we elicited muscle activity by stimulating the right sural nerve at 4–5xT to initiate local reflex and stimulating at a varying delay the left tibial or sural nerve to activate crossed reflex to investigate the interaction of the local and the crossed reflexes (**Fig. 5**). The delay between the two stimuli was chosen so that the center of the activity of the local reflex occurred in the silent period of the crossed reflex. We reasoned that if there is an inhibitory crossed reflex response, we should see a decreased EMG activity of the local reflex immediately after the left nerve stimulation (**Fig. 5a**, blue arrows). This inhibition was present in all recorded muscles regardless of whether the crossed reflex was initiated by the tibial nerve (**Suppl. Fig. 1**) or sural nerve stimulation (**Suppl. Fig 2**). This observation indicated the presence of an inhibitory crossed reflex response.

**Figure 5:**
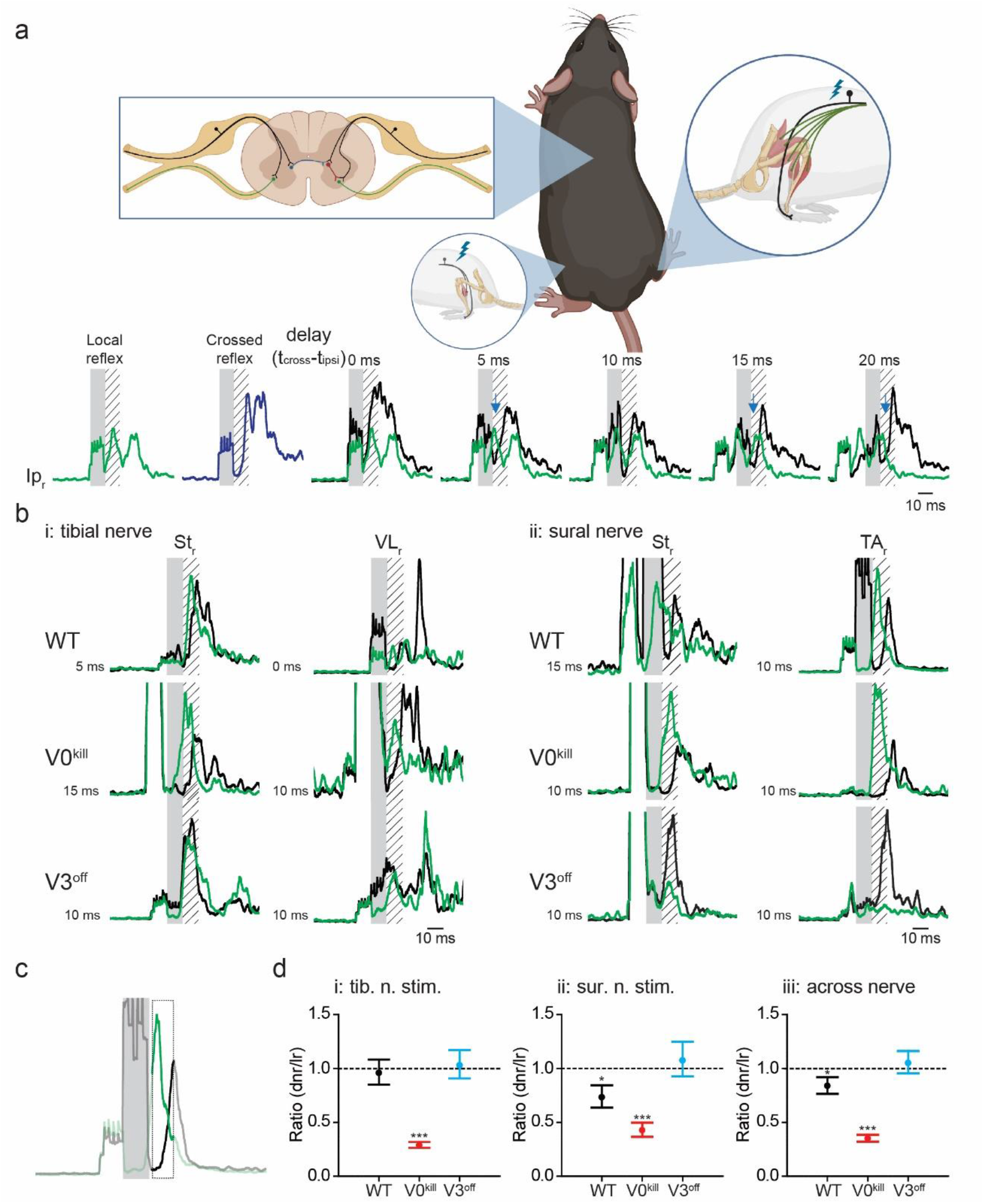
V3 CINs but not V0 CINs are necessary for the inhibitory crossed reflex. **a,** Schematic of the experimental setup. The experimental setup is as presented in **Fig. 1a** with the addition of one nerve stimulation electrode to the right sural nerve. The stimulation of the right sural nerve allowed us to elicit local reflex response (green trace) and stimulation of the left sural nerve the crossed reflex response (blue trace). The simultaneous stimulation of the left and right sural nerves with varying delays was used to detect inhibitory influence (black trace). Lower activity in black traces then green traces (blue arrows) indicated inhibitory crossed reflex action. Schematic was created with BioRender.com (agreement number: TZ250Q4IW4). **b,** Inhibitory effect was detected in the majority of EMG recordings in wild type and V0^kill^ mice but not in V3^off^ mice. **c,** To quantify the inhibitory crossed reflex, we averaged the green and black traces in the area indicated by the bold line (12 - 18 msec delay from stimulation onset) and took the ratio of these averages. Numbers smaller than 1 indicate inhibition. **d,** Graphs of ratios described in **c** for contralateral tibial (i) or sural (ii) nerve stimulation. The graph in iii is when the data in i and ii is pooled together. These graphs illustrate statistically, inhibition was preserved in V0^kill^ mice, but not in V3^off^ mice. dnr: reflex response to double nerve stimulation, lr: local reflex response. *: p<0.05, ***: p<0.001.

Interestingly, we observed that this inhibitory effect was preserved in V0^kill^ mice but was absent in V3^off^ mice (**Fig. 5b**). To confirm our observations quantitatively, we then activated the local reflex along with the crossed reflex, averaged the EMG activity within 12–18 msec time window from the onset of the left nerve stimulation and compared it to the average EMG activity of the local reflex only in the same time window (**Fig. 5c**). The ratio of these two parameters was used to quantify the amount of inhibition of the local reflex by the left nerve stimulation (values lower than 1 indicate inhibition). The generalized linear mixed model showed that there was a significant interaction effect between stimulation type (conditioning test vs. local reflexes only) and mouse type (WT, V3^off^, V0^kill^; F_2,2489_=149.659, p<0.0001), indicating that the amount of modulation of the local reflex by the crossed reflex in the 12–18 msec post-stimulation interval was different between the different mouse types (**Fig. 5d**). Post-hoc tests revealed that paired stimulation led to a significant reduction in the activity of the local reflex in WT (ratio=0.840±0.040, p=0.0002) and V0^kill^ mice (ratio=0.352±0.016, p<0.0001) but not in V3^off^ mice (ratio=1.053±0.053, p=0.306); and that the inhibition is significantly strongest in V0^kill^ mice, followed by WT and V3^off^ mice (all p≤0.031). Interestingly, we observed that the inhibition was more pronounced in V0^kill^ mice than in the wild type mice, suggesting that V0 CINs have a modulatory effect on crossed reflexes. Furthermore, a three-way interaction effect between stimulation type, mouse type and stimulated nerves (F_2,2244_=12.484, p<0.0001) suggests that the modulation of the local reflex activities by the crossed reflexes in WT, V3^off^, and V0^kill^ mice differed depending on whether sural or tibial nerve stimulation was applied. Indeed, post-hoc tests showed that the reduction of local reflex activities by the crossed reflexes in WT mice was only significant with sural nerve stimulation (ratio=0.733±0.052, p<0.0001) and not with tibial nerve stimulation (ratio=0.961±0.060, p=0.5255). In the case of V0^kill^ and V3^off^ mice, there were no differences between the nerves; the inhibition was present in V0^kill^ mice (sural: ratio=0.427±0.034, p<0.0001; tibial: ratio=0.290±0.014, p<0.0001) and absent in V3^off^ mice (sural: ratio=1.076±0.083, p=0.3434; tibial: ratio=1.031±0.067, p=0.6425) regardless of which nerve was stimulated. These data suggest the presence of the inhibitory crossed-reflex component in WT and V0^kill^ mice and that this inhibitory component was lost in V3^off^ mice.

Our data provide evidence that the inhibitory crossed reflex action elicited by tibial nerve or sural nerve stimulation is mediated by the exclusively excitatory V3 CINs. Interestingly, the V0 CINs which constitute excitatory and inhibitory CINs are not part of the inhibitory crossed reflex pathway, but can downregulate the inhibitory crossed reflex pathways.

### Involvement of V0 and V3 CINs in the crossed reflex actions during locomotion

We sought to provide additional evidence for the involvement of the V3 CINs in inhibitory crossed reflex pathways by investigating crossed reflex responses in the presence of muscle activity prior to reflex activation. Therefore, we investigated if the influence of crossed reflexes when muscles were activated by the premotor locomotor network prior to stimulation. To do this, we recorded the crossed reflex action during locomotion on a treadmill at a constant speed. All wild type mice were most comfortable moving at 0.2 m/s, therefore this speed was chosen for all wild type mice (**suppl. movie 1**). However, because the limited capability of locomotion is a main phenotypical characteristic of the V0^kill^ and V3^off^ mice, the locomotion for V3^off^ mice was set to 0.1 m/s, and V0^kill^ mice were set to 0.05 m/s (**suppl. movies 2, and 3, respectively**). It should be noted, that the V0^kill^ mice exhibited severe postural abnormalities making locomotion experiments especially challenging, but whenever they did stepping movements, the left and right hind limbs always stepped synchronously as previously described ^25^.

In accordance with our observation during the reflex response in non-locomoting animals, the data during locomotion provided evidence that V3 CINs are necessary for the inhibitory crossed reflex action, but the V0 CINs are not. To do this we visually inspected all cases in which the right sural (**Fig. 6a**) or tibial (**suppl. Fig. 3a**) nerve when there was activity in that particular muscle prior to nerve stimulation. We then calculated the occurrence of inhibition of the ongoing EMG activity as number of cases in which there was no activity in each left muscle within the 5 msec time window following the right sural (**Fig. 6b**) or the right tibial nerve (**suppl. Fig. 3b**) stimulation. In wild type and in V3^off^ mice, ongoing activity of muscles, except VL and TA, was suppressed by sural (**Fig. 6a and 6bi**) or tibial (**Suppl. Fig. 3a and 3bi**) nerve stimulation. The same was true for V0^kill^ mice when the tibial nerve was stimulated (**Suppl. Fig. 3bii**). Strikingly, during sural nerve stimulation, the inhibition was preserved in all muscles indicating the down-regulatory effect of inhibitory crossed reflex was diminished in the absence of V0 CINs (**Fig. 6bii**) suggesting that the downregulation of the inhibitory crossed reflex was compromised. Confirming the results presented in non-locomoting animals above, the termination of ongoing muscle activity after the left sural (**Fig6biii**) or tibial (**Suppl. Fig 3biii**) nerve stimulation was severely disrupted in V3^off^ mice, indicating that V3 CINs are necessary for the inhibitory crossed reflex also during locomotion. This observation was also confirmed when we quantified the temporal structure of the muscle activation pattern after the right sural (**Fig. 6c**) or tibial (**Suppl. Fig. 3c**) nerve stimulation. The onsets of muscle activity occurred on average around 10 msec after stimulation onset, sometimes even closer to zero indicating muscle activation as soon as stimulation started, in the absence of V3 CINs indicating no silent period after nerve stimulation (**Fig. 6d** and **Suppl. Fig. 3d**). In the absence of V0 CINs however, the onset latency was consistently closer to 20 msec in all recorded muscles, including the TA and VL, indicating the selective downregulation of inhibitory crossed reflex influence on TA and VL muscles was missing.

**Figure 6:**
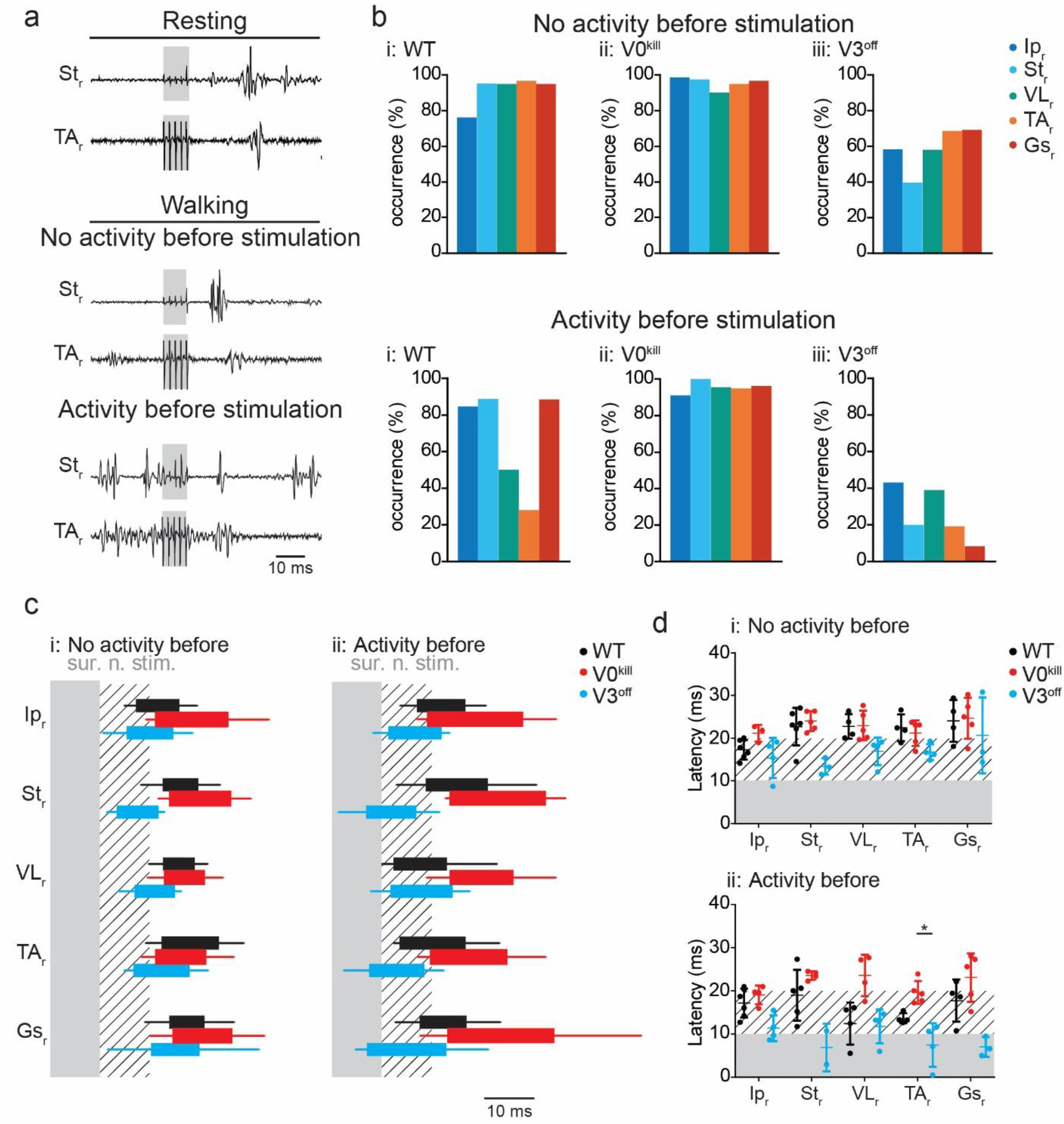
V3 CINs but not the V0 CINs are necessary for inhibitory crossed reflex during locomotion. **a,** Example EMG recordings from right St and TA muscles as a response to the left sural nerve stimulation during resting (top), and during locomotion when the muscle was either inactive (middle) or active (bottom) prior to the stimulation in a wild type mouse. **b,** Bar diagrams illustrating the probability of occurrence of a silent period in all recorded muscles of the right leg within the 5 msec window immediately after left sural nerve stimulation when there was no muscle activity (top) or there was activity (bottom) prior to nerve stimulation. **c,** Box diagrams showing the average (+/- standard deviation) on and offsets of right muscle activities as response to the left sural nerve stimulation at 5xT when there were no activity (i) or activity (ii) before stimulation (contralateral nerve stimulation indicated by the shaded area) during locomotion. Hatched area indicates 10 msec time window after nerve stimulation. Black bar: wild type, red bar: V0^kill^, and blue bar: V3^off^. **d,** Diagrams illustrating the onset latencies of muscle activities in Black bar: wild type (black), V0^kill^ (red), and V3^off^ (blue) mice during locomotion. Circles are averages from individual mice, and the horizontal lines indicate group averages (+/- standard deviation). *: p<0.05.

Together, we provide evidence that V3 CINs mediate the inhibitory crossed reflex response, and that V0 CINs are not involved in inhibitory crossed reflexes but are important for their downregulation during locomotion, as our data with resting animals suggested above.

## DISCUSSION

The current studies uncover the basic network structure that controls the excitatory and inhibitory crossed reflex pathways in mice. Our data suggest that the excitatory crossed reflex pathway is activated by proprioceptive group I afferents from muscle (1.2xT stimulation of the tibial nerve) as well as cutaneous afferent activation through sural nerve stimulation. In contrast, the inhibitory crossed reflex pathways were consistently activated by low threshold cutaneous afferents as indicated by 5xT stimulation of both nerves. We showed that the V0 CINs, which play a crucial role in left–right alternation of leg movement during locomotion, do not significantly contribute to the generation of the crossed reflexes but downregulate the inhibitory component. In contrast, we provided evidence that the V3 CINs contribute to the excitatory crossed reflex response and are an essential part of the inhibitory crossed reflex pathway.

The spinal circuitry that controls left–right coordination during locomotion is well understood in mice, due to advances in mouse genetics and electrophysiological techniques applicable to mice ^9^. Based on in vitro and in vivo experiments ^19,21,25,28^, a network design has been suggested that constitutes two inhibitory commissural pathways involving the V0 and the V3 CINs to coordinate left–right stepping in a speed-dependent manner ^26,28^. Despite this considerable advance in understanding the spinal locomotor circuitry, it is not known what the involvements of V0 and the V3 CINs are in the crossed reflexes. Previous observations that V0 CINs are necessary for maintaining left–right alternation in vivo ^25^ but also in vitro when the spinal cord is isolated and devoid of sensory afferent feedback ^19,20,25^ suggest that V0 CINs transmit information on the activity status of the CPG to the contralateral side. In contrast, although the pattern becomes more variable, the left–right alternating pattern is preserved in the absence of V3 CINs in vitro and in vivo ^21,28^. Our data present a reversed relationship when it comes to transmitting sensory information to the contralateral side. That is, V3 CINs seems to contribute to excitatory crossed reflex actions and are necessary for the inhibitory crossed reflex actions. In contrast, the V0 CINs are not directly involved in the crossed reflex pathway, however, they have a modulatory influence on the inhibitory crossed reflex actions. Therefore, our data provide evidence that previously defined CINs have distinct roles in involvement of crossed reflexes and regulating locomotor activities. A segregated commissural network for crossed reflexes and locomotion was previously suggested considering connectivity patterns of commissural interneurons with supraspinal centers and proprioceptive signals from muscles ^35^.

Our data suggest that V3 CINs are involved in the excitatory crossed reflex but crossed reflexes can still consistently be recorded when the V3 CINs are silenced. This can have two explanations. First, there are other main excitatory CINs involved in excitatory crossed reflex pathways, other than the V0 and the V3 CINs, such as glutamatergic excitatory CINs that derive from progenitor cells located dorsally in the embryonic spinal cord ^36^. Second, since silencing of the V3 CINs occurs during the embryonic development, the spinal circuitry might be reconfiguring so that other excitatory CINs, such as the V0_V_ might be compensating for the loss of V3 CIN functions. Our data do not allow to differentiate these two possibilities.

An interesting observation in our experiments was that the inhibitory crossed reflex actions transduced by cutaneous afferent signals were exclusively transmitted by excitatory CINs, the V3 CINs. This is in accordance with observations in cats that excitatory commissural interneurons synapse with local inhibitory interneurons, including the group Ia interneurons that mediate the reciprocal inhibitory stretch reflex response ^35^. A similar scheme was also suggested in humans ^18^ and in neonatal mice in vitro^37,38^. Our data provide evidence for a similar network design in mice regarding cutaneous afferent signals. That is, cutaneous sensory signals are conveyed by the exclusively excitatory V3 CINs to the contralateral side of the spinal cord to synapse with local inhibitory interneurons that in turn exert their inhibitory influence on the motor neurons. Support for this idea comes from previous histological experiments showing that the V3 CINs indeed synapse contralaterally with the local inhibitory interneurons including the V1 and V2b interneurons that among others, give rise to the group Ia interneurons and the Renshaw cells ^39,40^. Future experiments will have to reveal the identity of these local inhibitory interneurons that are involved in the inhibitory crossed reflex action. Nevertheless, our data add to the observation that inhibitory crossed reflex action is transmitted by excitatory commissural interneurons.

Crossed reflexes have been shown to be modulated during behavioral states or when the spinal circuitry is severed from the supraspinal centers ^7,8^. In our previous investigation, we have shown also in mice, the inhibitory crossed reflex actions are downregulated when the animal is moving ^31^. That confirms the previous observations in cats ^41^. However, the network underlying this modulation is not known. Our data provide evidence that the V3 CINs are the main CINs involved in the inhibitory crossed reflex pathways. The V0 CINs, however, are involved in the modulation of the inhibitory crossed reflex, based on three observations. First, inhibitory crossed reflex actions are much more pronounced in the absence of V0 CINs in V0^kill^ mice. Second, occurrence of the termination of an ongoing muscle activity during locomotion by contralateral nerve stimulation is much more pronounced in V0^kill^ mice. Third, more specifically, locomotion-dependent downregulation of the inhibitory crossed reflex selectively in VL and in TA muscles is absent during locomotion in V0^kill^ mice. The downregulation of the short latency inhibitory reflexes has been previously shown to be mediated by the brainstem ^42–44^ or by the leg position ^45^. Considering these observations, it is conceivable that the V0 CIN dependent downregulation of the inhibitory crossed reflex shown here, is a convergent with both mechanisms. The modulatory influence of V0 CINs on the inhibitory crossed reflex is indicated in **figure 7** as the dashed arrow from the V0 CINs to the inhibitory crossed reflex pathway.

**Figure 7:**
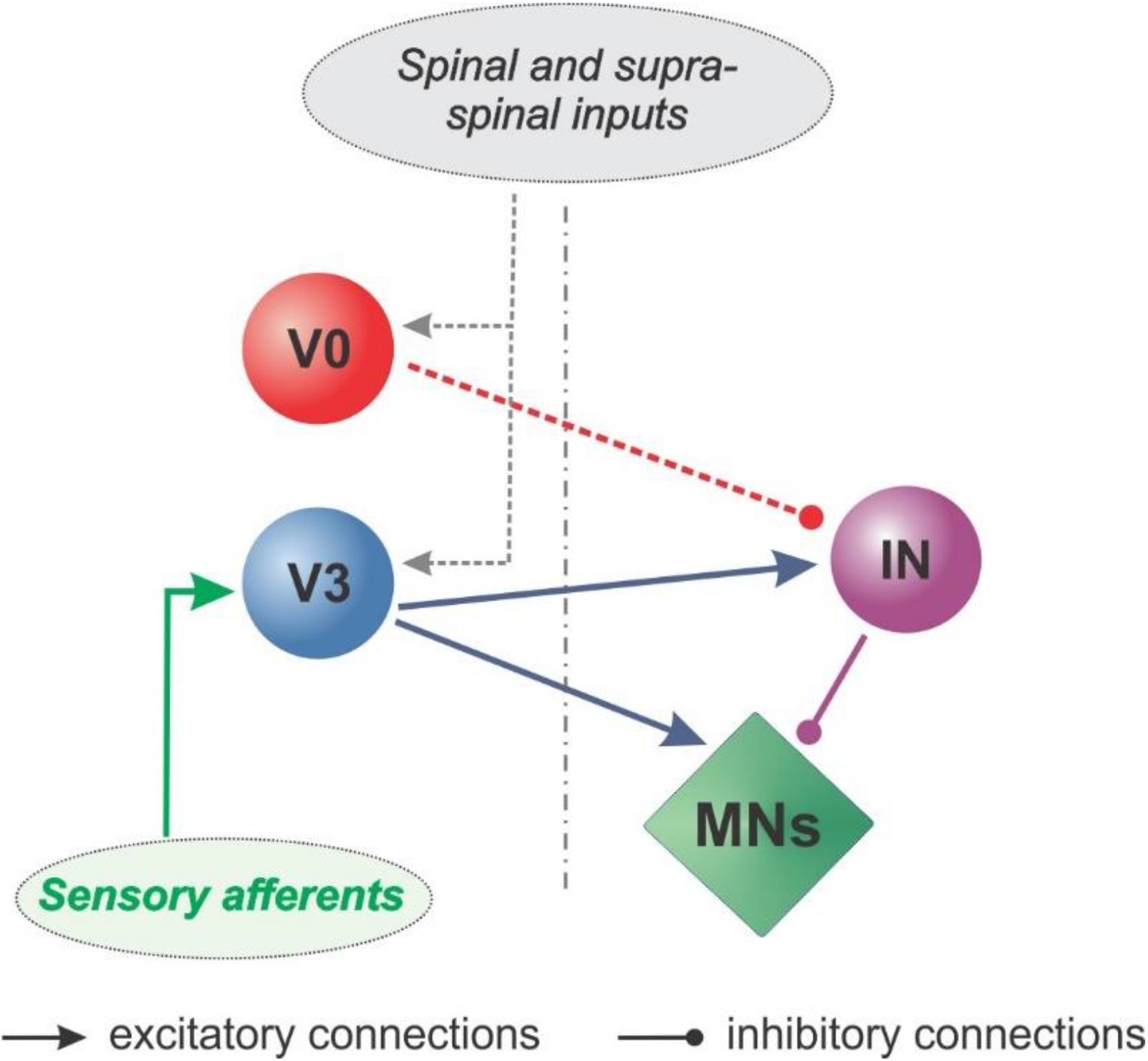
Commissural pathways involved in crossed reflexes. Our findings suggest that the spinal commissural pathways for crossed reflexes directly involve V3 CINs through a direct excitatory and an indirect inhibitory pathway. The inhibitory pathway is mediated at least with on local inhibitory interneuron (IN) as all V3 CINs are excitatory. V0 CINs although not directly involved in transmitting sensory afferent signals are modulate the inhibitory crossed reflex responses.

Our in vivo data are in accordance with previous observations that the presence of serotonin in neonatal spinal cord preparations in vitro downregulates synaptic transmission of inhibitory commissural interneurons with motor neurons ^37^. Given that serotonergic modulation is known to occur during locomotion ^46^, the downregulation of the inhibitory crossed reflexes that we observed could be due to the same mechanism. However, our findings extend the in vitro observation in two ways: first, we show this inhibitory commissural connection is possibly a part of the crossed reflex pathway. Second, the modulated inhibitory influence is limited to knee extensor (VL) and the ankle flexor (TA) muscles among the recorded muscles. These results provide the first insights into the identities of interneurons involved in the state-dependent modulation of the crossed reflex responses in vivo.

Our data provide insights into the spinal circuitry that controls crossed reflexes in mice and identifies genetically defined classes of commissural interneurons as part of it (**Fig. 7**). Our data do not identify any previously described subpopulations of the V3 CINs ^23,24^, but further research should shed light on which subpopulation of V3 CINs are the mediator of inhibitory crossed reflex. Similarly, our data do not provide information on the involvement of subpopulations of the V0 CINs in crossed reflexes. To identify which subpopulations of the V3 and the V0 CINs will be especially interesting as previous research has shown that they are differentially recruited in diverse behaviors ^27^. Future research should clarify these details and the modulation of the crossed reflex pathways during different motor behaviors.

Our data provide important insights into the crossed reflex circuitry as it identifies the involvement of the two major CINs, the V3 and the V0. We provide evidence that V3 CINs are directly involved in the crossed reflexes by transmitting sensory signals from one hind leg to the contralateral hind leg. V3 CINs are involved in the excitatory and the inhibitory crossed reflexes. As the V3 CINs are exclusively excitatory interneurons, there is at least one inhibitory local interneuron interconnected between the V3 CINs and the motor neurons. Furthermore, the V0 CINs although not directly involved, can downregulate inhibitory crossed reflexes. In addition, the V0 CINs exert a modulatory influence on the inhibitory crossed reflex action when the animal is moving. Our provides important detail into spinal circuitry for the crossed reflex pathways in the mouse, and further paves the way for further research into research to understand spinal circuitry for motor behavior.

## METHODS

### Animals

All experiments were performed according to the Canadian Council on Animal Care guidelines and approved by the Dalhousie University Committee on Laboratory Animals. Experiments were done on 19 wild type (15 male, 4 female), 19 V0^kill^ mice (11 male, 8 female), and 19 V3^off^ mice (7 male, 12 female). V0^kill^ mice (Hoxb8::Cre;Dbx1::DTA) were obtained by crossing the HoxB8::Cre mouse ^47^ provided as a courtesy of Dr. Hanns Ulrich Zeilhofer (University of Zurich) and the Dbx1::DTA mice obtained from Infrafrontier EMMA (Stock # EM 01926) as previously described ^25^. The V3^off^ mice (Sim1::Cre;Vglut2^flox/flox^) were obtained by crossing the Sim1::Cre mouse ^21^ with the Vglut2^flox/flox^ mouse ^48^ obtained from the Jackson Laboratories (Stock # 012898) as previously described ^24^. All mice were housed on a 12-hour light/dark cycle (light from 07:00 to 19:00) with access to laboratory chow and water ad libitum.

### Surgery

All adult mice (>6 weeks of age) were anesthetized with isoflurane (5% isoflurane with a constant flow rate of 1 l/min). Following the deep anesthesia was achieved indicated by a slow and regular breathing rate the anesthesia was maintained with 1.5–2% isoflurane throughout the surgery. At the onset of each surgery, ophthalmic eye ointment was applied to the eyes and the skin was sterilized by using a three-part skin scrub by means of Hibitane (Chlorhexidine gluconate 4%), alcohol, and povidone-iodine. At the beginning of the surgery buprenorphine (0.03 mg/kg) and ketoprofen (5 mg/kg) or meloxicam (5 mg/kg) were injected subcutaneously. A set of six bipolar EMG electrodes and one or two nerve stimulation cuffs were implanted in all experimental mice ^31^ as follows: small incisions were made on the shaved areas (neck and both hind limbs), and each bipolar EMG electrode and the nerve cuff electrodes were led under the skin from the neck incision to the leg incisions, and the headpiece connector was attached to the skin around the neck incision using suture. The EMG recording electrodes were implanted into the right hip flexor (iliopsoas, Ip_r_), knee flexor (semitendinosus, St_r_) and extensor (vastus lateralis, VL_r_), and ankle flexor (tibialis anterior, TA_r_) and extensor (gastrocnemius, Gs_r_), as well as the left ankle extensor (gastrocnemius, Gs_l_). Nerve stimulation electrodes were implanted in the left leg to activate contralateral proprioceptive and cutaneous feedback (tibial nerve) or predominantly cutaneous afferents (sural nerve), and the right leg to activate the right cutaneous afferents (sural nerve). After the incisions on the skin for electrode implantations were closed, anesthesia was discontinued, and mice were placed in a heated cage for at least 3 days before being returned to a regular mouse rack. Food mash and hydrogel were provided until full recovery after the surgery. Any handling of the mouse was avoided until the animal was fully recovered. The first recording session started at least 10 days after electrode implantation surgeries.

### Recording sessions

After the mice fully recovered from the implantation surgeries, the animals were briefly anesthetized with isoflurane, and a custom-made wire to connect the headpiece connector with the amplifier and the stimulation insulation units was attached to the mouse. The mice were removed from anesthesia and placed on a mouse treadmill (model 802; custom-built in the workshop of the Zoological Institute, University of Cologne, Germany). The electrodes were connected to an amplifier (model 102; custom-built in the workshop of the Zoological Institute, University of Cologne, Germany) and a stimulus isolation unit (ISO-FLEX; A.M.P.I., Jerusalem, Israel or DS4; Digitimer, Welwyn Garden City, UK). After the animal fully recovered from anesthesia (at least 5 min), we first determined the minimal (threshold) current necessary to initiate local reflex responses. To do this, we injected single impulses lasting 0.2 msec into the tibial nerve and double impulses each lasting 0.2 msec with 2 msec intervals into the sural nerve. The threshold currents were similar in wild type (tibial nerve: 110.6±45 mA; sural nerve: 448.1±416 mA), V0^kill^ (tibial nerve: 81.2±36 mA; sural nerve: 216.25±148.2 mA), and V3^off^ (tibial nerve: 143.1±113 mA; sural nerve: 236.7±135 mA) mice (P>0.05; Kruskal-Wallis with Dunn’s multiple comparison post-hoc test). These threshold currents then were used to set the current to either 1.2 times the threshold current (1.2xT, low current) or five times the threshold (5xT, high current) which was further used to elicit crossed reflex responses. These values were chosen because 1.2xT stimulation in mice activates group Ia and Ib afferents and 5xT stimulation activated in addition to group Ia and Ib afferents, also the group II muscle afferents from muscle spindles, and the group I cutaneous low-threshold mechanosensitive afferents ^32,33^.

Following the determination of threshold currents, the EMG signals from the five muscles of the right hind limb were recorded (sampling rate: 10 kHz for each muscle) while the right peripheral nerves, tibial nerve or the sural nerve, were stimulated at 1.2xT and 5xT with five brief impulses (impulse duration: 0.2 msec, frequency: 500 Hz) during resting (treadmill off and animal is either resting or calmly exploring the treadmill) or locomoting at a constant speed set by the treadmill. The treadmill was set at 0.2 m/s for all wild type mice as all wild type mice consistently locomoted at this speed (**Suppl. movie 1**). Because both mutant mice could not locomote steadily at higher speeds we set the treadmill speed for V3off mice at 0.1 m/s (**Suppl. movie 2**), and for V0kill mice at 0.05 m/s (**Suppl. movie 3**). In some experiments, the right sural nerve was also stimulated in combination with left tibial or sural nerve stimulation (wild type: 9 mice; V0^kill^: 18 mice; V3^off^: 12 mice). The EMG signals were amplified (gain 100), bandpass filtered, and stored on the computer using Power1410 interface and Spike2 software (Cambridge Electronic Design, Cambridge, UK).

### Statistics

Statistical analyses were performed in R 4.2.1 (R Foundation for Statistical Computing, Vienna, Austria). Generalized linear mixed models were calculated for various outcome variables and fit using Template Model Builder ^49^, interfaced through the glmmTMB R package^50^. Each model included a full factorial dispersion model to account for heteroskedasticity. Model assumptions were tested using the DHARMa package for R (distribution, dispersion, outliers, and quantile deviation tests were performed). Quantile–quantile (Q–Q) plots and histograms of residuals were inspected. Violation of assumptions led us to test different error distributions and link function. A Gamma-distribution with log-link function satisfied the assumptions of both models and was hence used. Pairwise post-hoc tests were performed for significant fixed effects and Tukey’s method was used to account for alpha-inflation due to multiple comparisons. An alpha-error of p<0.05 was regarded as significant.

Separate generalized linear mixed models with mouse line (WT, V3^off^, V0^kill^), stimulated nerve (tibial, sural), stimulation intensity (1.2x, 5x), and muscle, as well as their interaction effects as fixed effects and a per-animal random offset were run for 1) the average rectified EMG of muscle activity following nerve stimulation (**Fig. 2**), 2) the relative frequency of occurrence of muscle activity following nerve stimulation (**Fig. 3**), and 3) the onset latency of muscle activity (**Fig. 4**). For the average rectified EMG and onset latency, a Gamma-distribution with a log-link function and for the relative frequency of muscle activity a beta-distribution with logit-link function were used to ensure model assumptions were met.

To test if the inhibition of the local reflexes by the crossed reflexes differed between the wild type, V3^off^, and V0^kill^ mice, we calculated two parameters. First, we measured the average amplitude of the rectified and smoothened EMG traces within the 12–18 msec after the left nerve stimulation onset. Second, we measured the average EMG activity in the same time window when only the local reflex was activated (i.e., in the absence of contralateral nerve stimulation). That is, we measured the average of the rectified and smoothened EMG traces activated by the local reflex that would have corresponded to a 12-18 msec if the left nerve stimulation onset had occurred. To ensure reliable reflex recordings and avoid variability in data due to extended recording sessions, we developed exclusion criteria to only include recordings in which the reflex responses were stable through the experiments. Individual EMG recordings were excluded if: 1) there was persistent background activity (n=3 EMG recordings); 2) the crossed reflex (n=87 EMG recordings) was absent [defined as having an amplitude smaller than three times mean baseline activity (50 msec to 1 msec before the stimulation)]; 3) the local reflex was absent (n=11 EMG recordings); 4) the crossed (n=6 EMG recordings) or the local reflex (n=8 EMG recordings) were absent in the control recording after the protocol; and 5) finally, if either the crossed or local reflex size was larger than twice the size of the other reflex (n=27 EMG recordings). Thus, from a total of 212 recordings, 73 were included (WT: n=19, 11 sural, 8 tibial; V3^off^: n=23, 9 sural, 14 tibial; V0^kill^: n=31, 10 sural, 21 tibial). The stimulation type (paired nerve stimulation and local reflex only), stimulated nerve (sural and tibial nerve), and mouse line (WT, V3^off^, V0^kill^) as well as all second and third order interaction effects were modeled as fixed effects and a nested per mouse and muscle random offset was included. To account for non-normality, a Gamma-distribution with a log-link was used. To directly test whether the modulation of the local reflexes differed between mouse lines, contrasts comparing the effect of stimulation type (paired nerve stimulation vs local only) between each pair of mouse lines were calculated in addition to the regular post-hoc tests.

The graphical representations of data were made using GraphPad Prism 5 and processed using Illustrator CS5 (Adobe). All data are presented as means ± standard deviation. Normality was assess in GraphPad Prism 5 using omnibus K^2^ or Kolmogorov–Smirnov tests. ANOVA with Tukey post-hoc test when the data exhibited normal distribution or Kruskall-Wallis with Dunns comparison post-hoc test when data did not exhibit normal distribution was used to compare more than two groups. All statistical tests were two-tailed, and differences were considered statistically significant when the *P* value was <0.05.

## ACKNOWLEDGMENTS

The authors are grateful to Brenda Ross for maintaining the mouse colony and Dr. Kimberly Dougherty, Dr. Ilya Rybak, Dr. Rob Brownstone, and Dr. Sten Grillner for their comments on the manuscript. We also thank Dr. Hanns Ulrich Zeilhofer and Dr. Arthur Kania for providing the HoxB8::Cre breeder mice.

## AUTHOR CONTRIBUTIONS

Conceptualization: T.A.; Formal analysis: O.D.L, S.M.D, S.N.M; Experimentation: O.D.L., R.B.; Software: S.M.D, S.N.M; Providing Sim1::Cre mouse line: Y.Z.; Supervision: T.A.; Visualization: O.D.L, S.M.D, S.N.M; Writing original draft: T.A.; Review & editing: S.M.D., S.N.M. O.D.L., Y.Z., and T.A.

## Supplemental figures

**Suppl. Figure 1:**
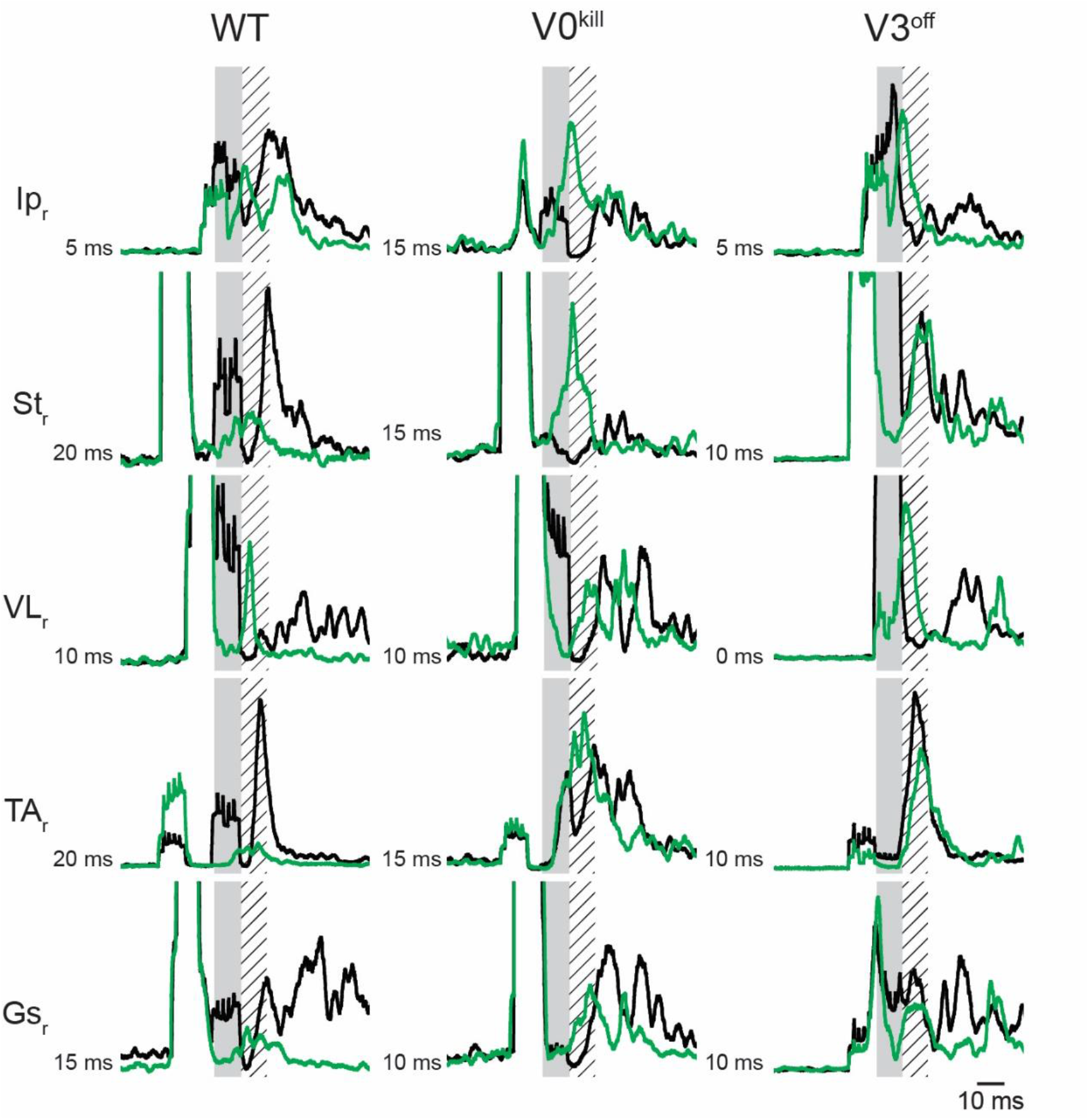
V3 CINs but not V0 CINs are necessary for the inhibitory crossed reflex initiated by the contralateral tibial nerve stimulation. Inhibitory effect was detected in the majority of all EMG recordings in wild type and V0^kill^ mice but not in V3^off^ mice, when the contralateral tibial nerve was stimulated.

**Suppl. Figure 2:**
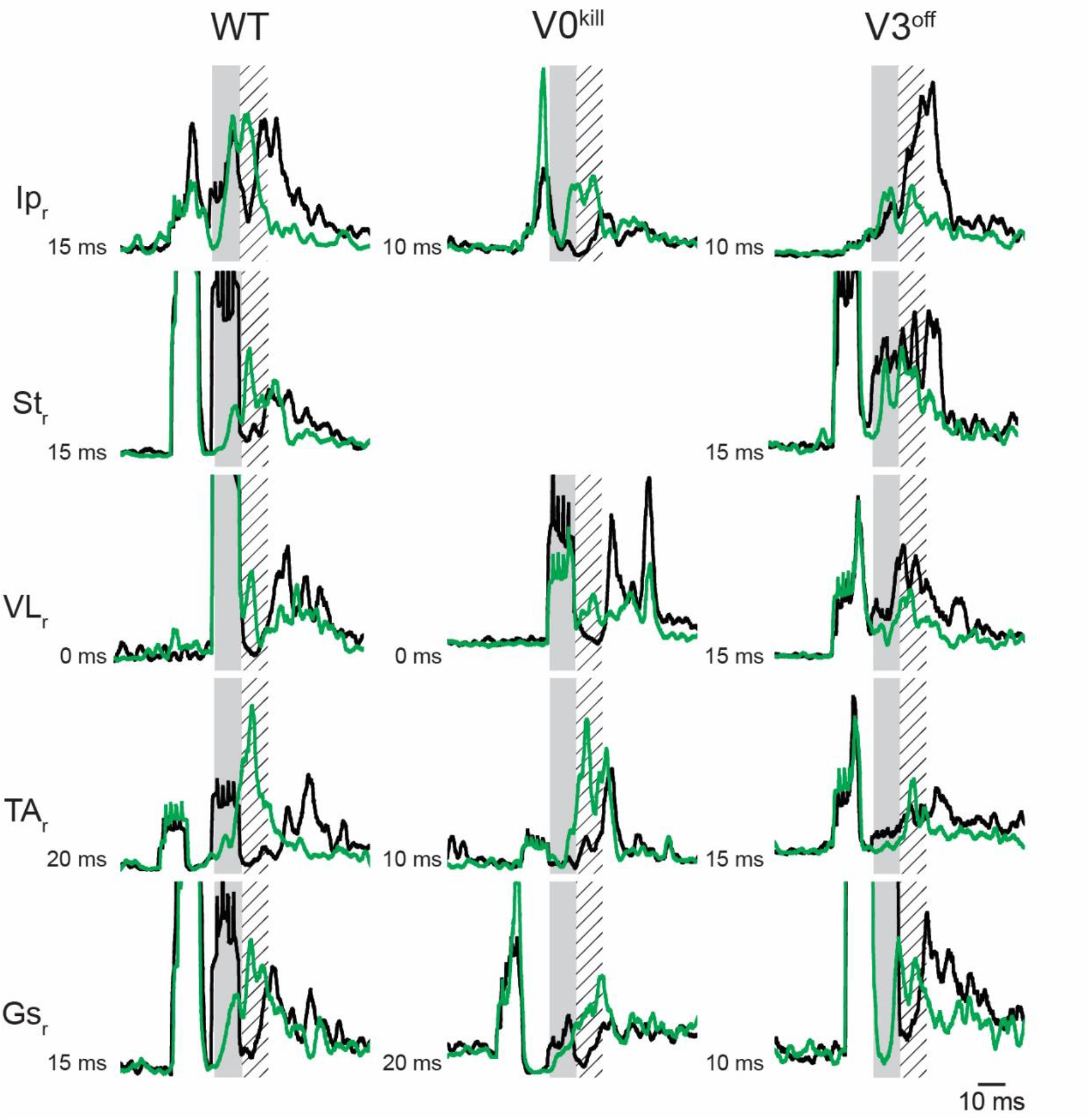
V3 CINs but not V0 CINs are necessary for the inhibitory crossed reflex initiated by the contralateral sural nerve stimulation. Inhibitory effect was detected in the majority of all EMG recordings in wild type and V0^kill^ mice but not in V3^off^ mice, when the contralateral sural nerve was stimulated.

**Suppl. Figure 3:**
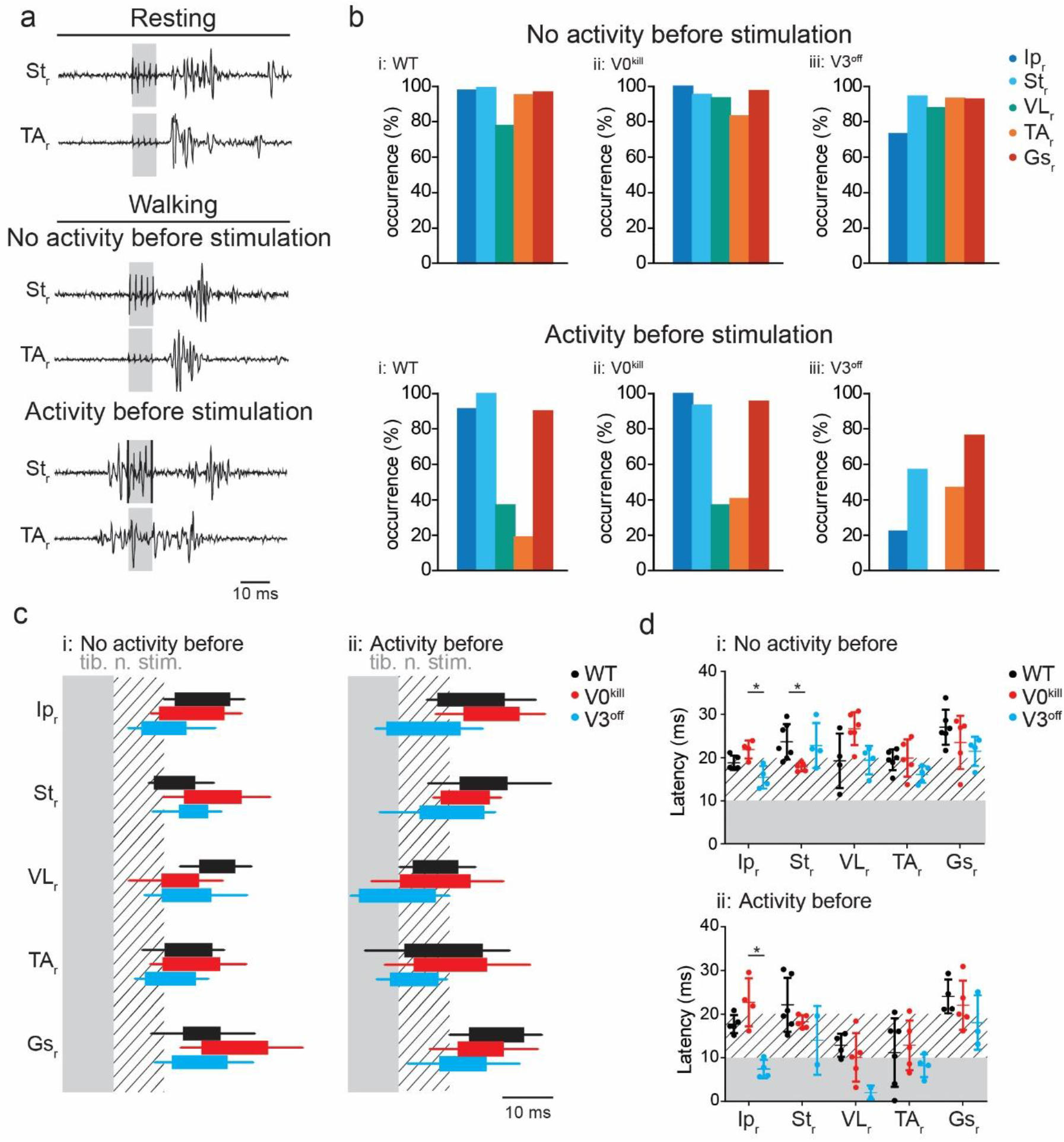
Crossed reflex responses to tibial nerve stimulation during locomotion. **a,** Example EMG recordings from right St and TA muscles as a response to the left tibial nerve stimulation during resting (top), and during locomotion when the muscle was either inactive (middle) or active (bottom) prior to the stimulation in a wild type mouse. **b,** Bar diagrams illustrating the probability of occurrence of a silent period in all recorded muscles of the right leg within the 5 msec window immediately after left tibial nerve stimulation when there was no muscle activity (top) or there were activity (bottom) prior to nerve stimulation. **c,** Box diagrams showing the average (+/- standard deviation) on and offsets of right muscle activities as response to the left tibial nerve stimulation at 5xT when there were no activity (i) or activity (ii) before stimulation (contralateral nerve stimulation indicated by the shaded area) during locomotion. Hatched area indicates 10 msec time window after nerve stimulation. Black bar: wild type, red bar: V0^kill^, and blue bar: V3^off^. **d,** Diagrams illustrating the onset latencies of muscle activities in Black bar: wild type (black), V0^kill^ (red), and V3^off^ (blue) mice during locomotion. Circles are averages from individual mice, and the horizontal lines indicate group averages (+/- standard deviation). *: p<0.05.

## REFERENCES

1. Jankowska, E. Spinal interneuronal networks in the cat: Elementary components. Brain Res. Rev. 57, 46–55 (2008).

2. Jankowska, E. & Edgley, S. A. Functional subdivision of feline spinal interneurons in reflex pathways from group Ib and II muscle afferents; an update. Eur. J. Neurosci. 32, 881–893 (2010).

3. Arya, T., Bajwa, S. & Edgley, S. A. Crossed reflex actions from group II muscle afferents in the lumbar spinal cord of the anaesthetized cat. J. Physiol. 444, 117–131 (1991).

4. Edgley, S. A., Jankowska, E., Krutki, P. & Hammar, I. Both dorsal horn and lamina VIII interneurones contribute to crossed reflexes from feline group II muscle afferents. J. Physiol. 552, 961–974 (2003).

5. Sherrington, C. S. On Reciprocal Innervation of Antagonistic Msucles.-Eighth Note. Proc. R. Soc. B 76, 269–297 (1905).

6. Gauthier, L. & Rossignol, S. Contralateral hindlimb responses to cutaneous stimulation during locomotion in high decerebrate cats. Brain Res. 207, 303–320 (1981).

7. Perl, E. R. Crossed Reflexes of Cutaneous Origin. Am. J. Physiol. 188, 609–615 (1957).

8. Duysens, J., Loeb, G. E. & Weston, B. J. Crossed flexor reflex responses and their reversal in freely walking cats. Brain Res. 197, 538–542 (1980).

9. Kiehn, O. Decoding the organization of spinal circuits that control locomotion. Nat. Rev. Neurosci. 17, 224–238 (2016).

10. Jankowska, E. Spinal interneurons. in Neuroscience in the 21st Century: From Basic to Clinical (ed. Pfaff, D. W.) 11063–1099 (Springer Science+Business Media, 2013). doi:10.1007/978-1-4614-1997-6

11. Jankowska, E. Spinal Reflexes. in Neuroscience in the 21st Century (eds. Pfaff, D. W. & Volkow, N. D.) 1599–1621 (Springer Science+Business Media, 2016). doi:10.1007/978-1-4614-1997-6

12. Dietz, V., Colombo, G., Jensen, L. & Baumgartner, L. Locomotor capacity of spinal cord in paraplegic patients. Ann. Neurol. 37, 574–582 (1995).

13. Plotnik, M., Giladi, N. & Hausdorff, J. M. A new measure for quantifying the bilateral coordination of human gait: effects of aging and Parkinson’s disease. Exp. Brain Res. 181, 561–570 (2007).

14. Tseng, S. & Morton, S. M. Impaired Interlimb Coordination of Voluntary Leg Movements in Poststroke Hemiparesis. J. Neurophysiolgy 104, 248–257 (2010).

15. Meijer, R. et al. Markedly impaired bilateral coordination of gait in post-stroke patients : Is this deficit distinct from asymmetry ? A cohort study. J. Neuroeng. Rehabil. 8, 23 (2011).

16. Krasovsky, T. et al. Stability of gait and interlimb coordination in older adults. J. Neurophysiol. 107, 2560–2569 (2012).

17. Stubbs, P. W., Nielsen, J. F., Sinkjær, T. & Mrachacz-Kersting, N. Short-latency crossed spinal responses are impaired differently in sub-acute and chronic stroke patients. Clin. Neurophysiol. 123, 541–549 (2012).

18. Mrachacz-Kersting, N., Geertsen, S. S., Stevenson, A. J. T. & Nielsen, J. B. Convergence of ipsi- and contralateral muscle afferents on common interneurons mediating reciprocal inhibition of ankle plantar flexors in humans. Exp. Brain Res. 235, 1555–1564 (2017).

19. Bellardita, C. & Kiehn, O. Phenotypic Characterization of Speed-Associated Gait Changes in Mice Reveals Modular Organization of Locomotor Networks. Curr. Biol. 25, 1426–1436 (2015).

20. Lanuza, G. M., Gosgnach, S., Pierani, A., Jessell, T. M. & Goulding, M. Genetic Identification of Spinal Interneurons that Coordinate Left-Right Locomotor Activity Necessary for Walking Movements. Neuron 42, 375–386 (2004).

21. Zhang, Y. et al. V3 Spinal Neurons Establish a Robust and Balanced Locomotor Rhythm during Walking. Neuron 60, 84–96 (2008).

22. Pierani, A. et al. Control of Interneuron Fate in the Developing Spinal Cord by the Progenitor Homeodomain Protein Dbx1. Neuron 29, 367–384 (2001).

23. Borowska, J. et al. Functional Subpopulations of V3 Interneurons in the Mature Mouse Spinal Cord. J. Neurosci. 33, 18553–18565 (2013).

24. Chopek, J. W., Nascimento, F., Beato, M., Brownstone, R. M. & Zhang, Y. Sub-populations of Spinal V3 Interneurons Form Focal Modules of Layered Pre-motor Microcircuits. Cell Rep. 25, 146–156.e3 (2018).

25. Talpalar, A. E. et al. Dual-mode operation of neuronal networks involved in left-right alternation. Nature 500, 85–88 (2013).

26. Shevtsova, N. A. et al. Organization of left – right coordination of neuronal activity in the mammalian spinal cord : Insights from computational modelling Key points. J. Physiol. 11, 2403–2426 (2015).

27. Zelenin, P. V, Vemula, M. G., Lyalka, V. F., Kiehn, O. & Deliagina, T. G. Differential Contribution of V0 Interneurons to Execution of Rhythmic and Nonrhythmic Motor Behaviors. J. Neurosci. 41, 3432–3445 (2021).

28. Zhang, H. et al. The role of V3 neurons in speed-dependent interlimb coordination during locomotion in mice. Elife 11, e73424 (2022).

29. Danner, S. M., Wilshin, S. D., Shevtsova, N. A. & Rybak, I. A. Central control of interlimb coordination and speed-dependent gait expression in quadrupeds. J. Physiol. 594.23, 6947–6967 (2016).

30. Danner, S. M., Shevtsova, N. A., Frigon, A. & Rybak, I. A. Computational modeling of spinal circuits controlling limb coordination and gaits in quadrupeds. Elife 6, e31050 (2017).

31. Laflamme, O. D. & Akay, T. Excitatory and inhibitory crossed reflex pathways in mice. J. Neurophysiol. 120, 2897–2907 (2018).

32. Schomburg, E. D., Kalezic, I., Dibaj, P. & Steffens, H. Reflex transmission to lumbar α-motoneurones in the mouse similar and different to those in the cat. Neurosci. Res. 76, 133–140 (2013).

33. Steffens, H., Dibaj, P. & Schomburg, E. D. In Vivo Measurement of Conduction Velocities in Afferent and Efferent Nerve Fibre Groups in Mice. Physiol. Res. 61, 203–214 (2012).

34. Peyronnard, J.-M. & Charron, L. Motor and sensory neurons of the rat sural nerve: A horseradish peroxide study. Muscle Nerve 5, 664–660 (1982).

35. Jankowska, E., Krutki, P. & Matsuyama, K. Relative contribution of Ia inhibitory interneurones to inhibition of feline contralateral motoneurones evoked via commissural interneurones. J. Physiol. 568.2, 617–628 (2005).

36. Alaynick, W. A., Jessell, T. M. & Pfaff, S. L. Snapshot : Spinal Cord Development. Cell 146, 178–178.e1 (2011).

37. Butt, S. J. B. & Kiehn, O. Functional Identification of Interneurons Responsible for Left-Right Coordination of Hindlimbs in Mammals. Neuron 38, 953–963 (2003).

38. Quinlan, K. A. & Kiehn, O. Segmental, Synaptic Actions of Commissural Interneurons in the Mouse Spinal Cord. J. Neurosci. 27, 6521–6530 (2007).

39. Zhang, J. et al. V1 and V2b Interneurons Secure the Alternating Flexor-Extensor Motor Activity Mice Require for Limbed Locomotion. Neuron 82, 138–150 (2014).

40. Alvarez, F. J. et al. Postnatal phenotype and localization of spinal cord V1 derived interneurons. J Comp Neurol 493, 177–192 (2005).

41. Frigon, A. & Rossignol, S. Short-latency crossed inhibitory responses in extensor muscles during locomotion in the cat. J. Neurophysiol. 99, 989–998 (2008).

42. Eccles, R. M. & Lundberg, A. Supraspinal control of interneurones mediating spinal reflexes. J. Physiol. 147, 565–584 (1959).

43. Holmqvist, B. & Lundberg, A. Differential supraspinal control of synaptic actions evoked by volleys in the flexion reflex afferents in alpha motoneurons. Acta Physiol. Scand. 186, 1–15 (1961).

44. Grillner, S. & Shik, M. L. On the Descending Control of the Lumbosacral Spinal Cord from the “ Mesencephalic Locomotor Region Acta Physiolgica Scand. 87, 320–333 (1973).

45. Grillner, S. & Rossignol, S. Contralateral reflex reversal controlled by limb position in the acute spinal cat injected with clonidine i.v. Brain Res. 144, 411–414 (1978).

46. Jordan, L. M., Liu, J., Hedlund, P. B., Akay, T. & Pearson, K. G. Descending command systems for the initiation of locomotion in mammals. Brain Res. Rev. 57, 183–191 (2008).

47. Witschi, R. et al. Hoxb8-Cre mice: A tool for brain-sparing conditional gene deletion. Genesis 48, 596–602 (2010).

48. Tong, Q. et al. Synaptic Glutamate Release by Ventromedial Hypothalamic Neurons Is Part of the Neurocircuitry that Prevents Hypoglycemia. Cell Metab. 5, 383–393 (2007).

49. Kristensen, K., Nielsen, A., Berg, C. W., Skaug, H. & Bell, B. M. TMB: Automatic Differentiation and Laplace Approximation. J. Stat. Softw. 70, 1–21 (2016).

50. Brooks, M. E. et al. glmmTMB Balances Speed and Flexibility Among Packages for Zero-inflated Generalized Linear Mixed Modeling. R J. 9, 378–400 (2017).

